# Stratification and prediction of drug synergy based on target functional similarity

**DOI:** 10.1101/586123

**Authors:** Mi Yang, Michael P. Menden, Patricia Jaaks, Jonathan Dry, Mathew Garnett, Julio Saez-Rodriguez

## Abstract

Targeted mono-therapies in cancer are hampered by the ability of tumor cells to escape inhibition through rewiring or alternative pathways. Drug combination approaches can provide a means to overcome these resistance mechanisms. Effective use of combinations requires strategies to select combinations from the enormous space of combinations, and to stratify patients according to their likelihood to respond. We here introduce two complementary workflows: One prioritising experiments in high-throughput screens for drug synergy enrichment, and a consecutive workflow to predict hypothesis-driven synergy stratification. Both approaches only need data of efficacy of single drugs. They rely on the notion of target functional similarity between two target proteins. This notion reflects how similarly effective drugs are on different cancer cells as a function of cancer signaling pathways’ activities on those cells. Our synergy prediction workflow revealed that two drugs targeting either the same or functionally opposite pathways are more likely to be synergistic. This enables experimental prioritisation in high-throughput screens and supports the notion that synergy can be achieved by either redundant pathway inhibition or targeting independent compensatory mechanisms. We tested the synergy stratification workflow on seven target protein pairs (AKT/EGFR, AKT/MTOR, BCL2/MTOR, EGFR/MTOR, AKT/BCL2, AKT/ALK and AKT/PARP1, representing 29 combinations and predicted their synergies in 33 breast cancer cell lines (Pearson’s correlation r=0.27). Additionally, we experimentally validated predicted synergy of the BRAF/Insulin Receptor combination (Dabrafenib/BMS−754807) in 48 colorectal cancer cell lines (r=0.5). In conclusion, our synergy prediction workflow can support compound prioritization in large scale drug screenings, and our synergy stratification workflow can select where the efficacy of drugs already known for inducing synergy is higher.

## INTRODUCTION

In the quest for clinical efficacy, drug combinations are a promising strategy in cancer treatment(1,2). Targeting a signaling pathway at one step may not be sufficient for reaching maximal effects on pathway inhibition. Using one agent at higher dose could be a short term solution. However, higher dose leads to increased toxicity and emergence of resistance to treatments. Resistance mechanisms to monotherapy can occur by activation of compensatory signaling. For example, the activation of ERK signaling in melanoma when treated with BRAF inhibitors may lead to paradoxical activation of CRAF(3). Targeting BRAF and downstream MEK at the same time proved to be beneficial for overall patient survival(4), by inhibiting the initial BRAF driver mutation and paradox CRAF activation. Alternatively to inhibiting two key proteins within the same pathway, a common strategy is to parallel inhibit two separate cancer pathways to maximise drug efficacy. For example, parallel inhibition of ERK and AKT could be beneficial as those pathways may be connected through cross talks and feedback loops in breast cancer(5). Given the enormous space of potential drug combinations, strategies to effectively predict their efficacy are highly desirable.

Many methods predict drug synergy using chemical structure and genomic information(6–8). Preuer *et al.* used deep learning to predict synergy within the space of explored drugs and cell lines (Pearson’s correlation of observed versus predicted synergy score r=0.73), but observed a much worse performance in predicting untested drugs (r=0.48) or untested cell lines (r=0.57)(8). Jaeger *et al* identified new drug combinations using network topology of pathway cross-talk(9). However, gene mutation information, arguably the most actionable information in the clinic, was not used. In the recent Dialogue on Reverse-Engineering Assessment and Methods (DREAM) drug combination challenge(10), the best performing team used a protein-protein interaction network to augment the genomic features based on their network distance from drug targets. Whilst the best performer achieved outstanding predictability comparable to the level of experimental replicates, synergy was predicted based on supervised machine learning algorithms. A common bottleneck for the application of all supervised learning methods is the limited publicly available combinatorial drug screening data. In practice, the combinatorial explosion of drug pairs is the limiting factor to both the number of experimentally tested drugs, and the number of tested cell lines. Additionally, tested combinations are driven by expert’s knowledge, and therefore may be focused on known biological examples and thereby bias the performance of supervised learning.

Synergy of combination can be estimated from the effect of single drugs. For example, in the NCI-DREAM Drug Sensitivity and Drug Synergy Challenge(7) the similarities on the effect of drugs on gene expression was used to predict synergy. However, this requires the generation of expression data upon treatment, which is relatively costly.

We here investigate if we can use similarities of single drugs in just the effect on cell survival to learn about the efficacy of combinations. We propose a methodology for prioritizing drug combinations and for cell line stratification based on the functional similarity between two target proteins. For this, we extend the notion of compound similarity to target similarity: the functional similarity of a pair of target proteins is defined as the correlation between the drug response upon perturbation of those proteins, as a function of the activity of a set of essential pathways. Pathway activities are computed from data-derived gene sets, that have been demonstrated to be more predictive than pathway-based gene sets(11,12). Different cancer types may be driven by different cancer pathways. Therefore, the similarity metric is context dependent. Two target proteins that are functionally very similar are likely to belong to the same signaling pathway; on the contrary, functionally dissimilar proteins are likely to belong to unrelated pathways. We find higher synergy likelihood when there is either very high or very low similarity. Based on this information, we build a compound prioritization methodology for high-throughput screens. Furthermore, we explore context specific (breast and colorectal cancer) drug combinations for their mode of actions based on known mechanistic insights from the monotherapies, to predict synergy and potentially enable patient stratifications in the clinic.

## RESULTS

### Synergy prediction workflow

#### Profiling Target functional similarity with matrix factorization

We defined the similarity in drug response profile with respect to a set of essential pathways when targeting two proteins. We used the Genomics of Drug Sensitivity in Cancer (GDSC) data(13) composed of 990 cancer cell lines, treated by 265 drugs, with deep molecular characterization of the cell lines including gene expression, methylation, DNA mutation and copy number variation profiles.

We first computed the pathway activity from gene expression of the GDSC data using Pathway RespOnsive GENes (PROGENy(12), **Methods, Fig. 1a**). Next, we applied the Macau algorithm(14) to find interactions between the drugs’ nominal targets and pathway activities(15) (**Methods, Fig. 1a**). We considered two target proteins, each targeted by a different drug, and took the Pearson’s correlation between their interactions with all PROGENy pathways. We defined this correlation as the functional similarity of those two proteins. A pathway contains more information than a single gene’s expression level. Therefore, functional similarity based on a small subset of essential pathways is likely to be more robust than using thousands of genes, of which the vast majority are not involved in drug response or in cancer.

**Fig. 1:**
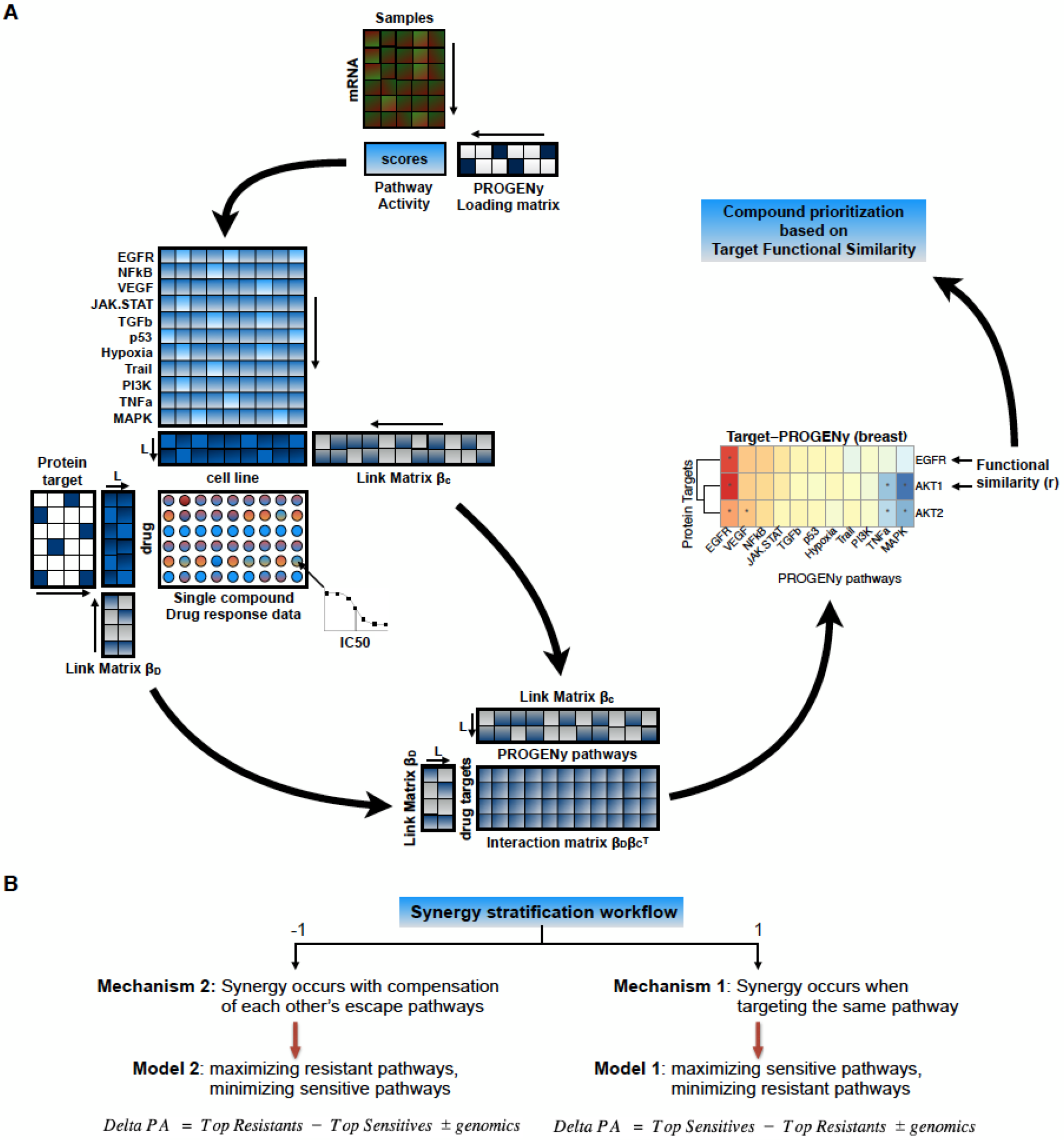
Methodology for drug synergy prediction and stratification. **A**. (i) First, we compute activities scores for 11 pathways from gene expression of cancer cell lines. It consists in multiplying the transcriptomics data by a loading matrix using PROGENy(12). (ii) We then use Macau algorithm(14) to predict multiple drugs’ responses simultaneously by uncovering the common (latent) features that can benefit each individual learning task. We use the previously derived pathway scores as input features (side information) for cell lines, and nominal target for drugs. Each side information matrix is transformed into a matrix of L latent dimensions by a link matrix. Drug response is then computed by a matrix multiplication of the two latent matrices. (iii) Concurrently to drug response prediction, we derive the interactions between drug features (targets) and cell line features (pathway activity), by multiplying the two link matrices. An association between protein X and pathway Y means that activation of pathway Y correlates with drug sensitivity when targeting protein X. In case of causality, we can say that activation of pathway Y confers sensitivity to any drug targeting protein X. (iv) These interactions allow us to define the functional similarity between two target proteins. In this example of breast tissue, The functional similarity between proteins EGFR and AKT1 is the correlation of their interaction values with the 11 PROGENy pathways. As final step of the synergy prediction workflow, the derived target functional similarity informs us about the likelihood of synergy. (v) We use the Target functional similarity metric for compound prioritization. **B**. For synergy stratification workflow, we start with target pairs already known to be synergistic. The value of the functional similarity between the target proteins reflects different synergy mechanisms. If the similarity is close to 1, synergy occurs by targeting the same signaling pathways. A similarity close to −1 suggests a synergy induced by compensation of escape mechanism. We build specific synergy model for each case to predict synergy scores of cancer cell lines.

#### Functional similarity of drugs’ targets influences drug synergy

To answer the question whether the functional similarity of drugs’ targets affect synergy, we used the AstraZeneca drug combination DREAM challenge data(10), composed of 910 combinations and 85 cancer cell lines, which are also part of the GDSC panel. We selected the 25 target proteins from GDSC that are also part of the DREAM challenge data (**Fig. 2a**). There are 300 pairwise combinations from the n=25 proteins, from which we selected 99 pairs where the two proteins are targeted by two different drugs in the GDSC panel, since we are interested in drug combinations. We considered the target pathway interaction matrix and for each combination of targets, we computed the Pearson’s correlation of the interaction score with the PROGENy pathways. The target combinations were then ranked from the most correlated pair to the most anti-correlated pair. For instance, the proteins BRAF and MEK are in the same pathway (ERK signalling), have a functional similarity of 0.74 (P=0.0088) in skin cell lines, and are synergistic within this cancer type(16). We consider that if the similarity between two target proteins is greater than 0.7 (**Supplementary Fig. 7)**, then inhibition of the proteins triggers similar effects.

**Fig. 2:**
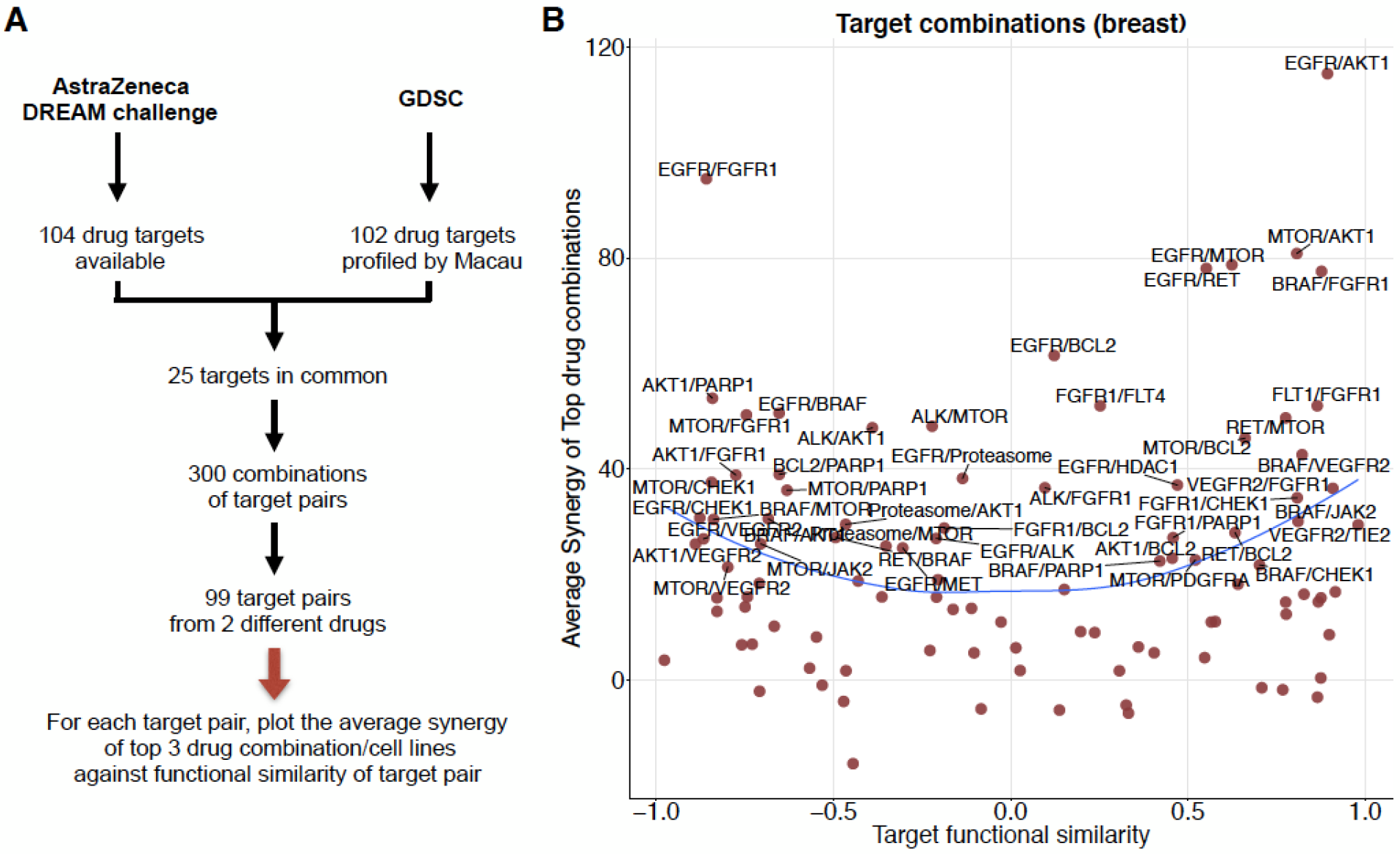
Influence of the similarity between target proteins on drug synergy. **A.** We selected common targets from AstraZeneca and GDSC data sets. **B.**The target functional similarity is the correlation between 2 targets by their interactions with the PROGENy pathways. A correlation of 1 implies that the activities of pathways correlates in the same way with drug efficacy on those proteins. A correlation of −1 implies opposite effects. The average synergy is computed for each target pair, as the mean of the top three synergistic drug-cell line pairs. We chose a threshold of 20 as synergistic effect, and a score lower than −20 as antagonistic effect, as in Menden *et al(10)*.

Synergy scores in AstraZeneca dataset are derived from the volume difference of an experimental 5-by-5 matrix (combinations tested at different drug concentrations) and the theoretical Loewe additivity surface inferred from both monotherapies (**Methods**). To ascertain if target functional similarity can influence drug synergy, we chose the breast tissue as it is the most represented in the dataset (33 cell lines), and for each target pair, we plotted the observed average synergy scores of the top three synergistic drug1-drug2-cell triplets, against the target functional similarity of this target pair (**Fig. 2b**). We observed that synergy arises in both highly correlated and highly anti-correlated target groups (**Fig. 2b**). Remarkably, very few synergistic target pairs were found with a functional similarity close to zero (lowly correlated target group).

We tested the significance of our observation on breast (33 cell lines), colon (12 cell lines) and NSCLC (22 cell lines), by predicting the average observed synergy score of the top synergistic combinations using the absolute value of the target functional similarity (**Supplementary Fig. 1**). For breast, colon and lung tissues, the prediction performances (Pearson’s correlation) are r=0.25 (P=0.014), r=0.45 (P=0.27) and r=0.14 (P=0.56), respectively. The trend is stronger for colon compared to lung, which is why we chose to focus on colorectal cancer cell lines.

#### External validation of the functional similarity metric

We further validated those trends on the NCI-ALMANAC dataset(17). We used functional similarity from Sanger data to predict the synergy in NCI-ALMANAC. For 18 target pairs and 8 tissues in common, the prediction performance is r=0.35 (P=0.15) (**Supplementary Fig. 2a**), using the average functional similarity across all tissues to predict the average synergy. The small number of targets (18 versus 99 in AstraZeneca data) makes the average across tissues potentially more reliable than taking individual tissues. Although the performance is exceptional for brain and colon tissues, r=0.75 (P=0.0022) and r=0.32 (P=0.25), respectively (**Supplementary Fig. 2c, d**), and failed for breast, r=-0.33 (P=0.025).

These results made us hypothesize that target functional similarity based on pathway activations is a metric that can be used for compound prioritization: for any given target pair, the more functional similar or opposite two proteins are, the most likely synergy will arise. We reason that this could be due to complementary mechanisms of synergy that take place: **Mechanism 1 (Synergy by similarity)**: When two drugs have similar interaction profiles, they are most likely targeting some common mechanism. In this case, synergy may be achieved by double hit of the same pathway, or putatively inhibiting feedback loops. **Mechanism 2 (Synergy by compensation)**: In contrast, for functionally opposite proteins, when one pathway’s activation is correlated with drug sensitivity for targeting one protein, it is also correlated with resistance for targeting the other protein. This functional landscape may prevent the activation of compensatory escape pathways, and thereby increases the likelihood to observe synergy.

We developed a workflow for ranking synergy enrichment (**Fig. 1a**), based on multitask learning through the following steps: (i) Compute tissue specific interaction matrix between target proteins and pathway activities using single drug screening data. (ii) Find common target proteins between single drug screening data and drug synergy data. (iii) Compute pairwise Target functional similarity in single drug screening data for the selected common targets. (iv) Keep target pairs with absolute Target functional similarity greater than a certain threshold (for example: 0.7). Our method returns a ranking of experimentally untested drug combinations from being likely to unlikely synergistic, which ultimately enables a prioritization for future experiments.

### Synergy stratification workflow

#### Stratifying cancer cell lines for synergistic combinations

As an addition to the synergy prediction workflow, we propose a consecutive step to stratify cell lines from responders to non responders. For this, we use the inferred synergy mechanism and pathway activities of new samples to build specific models to predict synergy for new drug combinations. The synergy stratification workflow predicts the actual synergy scores on samples for a given target pair for which synergy has been described (either through experiments or from literature). For each of the previously described synergy mechanisms, we built specific models to predict synergy scores on new cancer cell lines (**Methods, Fig. 1b**). We used Delta Pathway Activity (Delta PA), a linear combination of pathway activity, SNP and CNV to predict synergy. Therefore, our models stratify the cell lines based on their “genomic context” (**Methods**). We only considered drug combination known to induce synergy, for the following reasons: (i) In practice, we decide about stratification only after knowledge of synergy potential. (ii) We predict synergy score with a linear combination of pathway activities (**Methods**), a relative concept that is not based on actual synergy scores, and therefore not designed to predict the existence of synergy.

We propose a general framework to predict synergy scores follows several key steps and we emphasize on the notion of “target combination” which represents the dual inhibition of two target proteins, regardless of the drugs that are used (**Methods**). We applied our methodology to AstraZeneca drug combination data for breast tissue and experimentally validated a predicted synergistic drug combination for colorectal cancer cell lines.

#### Application to the AstraZeneca breast data set

We tested our synergy models on different target pairs by computing the Pearson’s correlation of observed versus predicted synergy scores on all available cell lines. The observed synergy is computed as the average of all drug combinations targeting a given target pair, across all available cell lines. Therefore, for each cell line, the observed average synergy may be computed for different drug combinations since the matrix of drug-cell line synergy is sparse.

We selected target pairs that fulfilled the following conditions: 1) Observed CombeneFit synergy score(18) of top hits must be greater than 20, considered as a clear threshold for synergy(10)(**Fig. 2b**). 2) Drug combinations have had to be tested in at least 10 cell lines, owing to the limitations of measuring performance by Pearson’s correlation. 3) At least two different drug combinations for the target pair were tested in each cell line, otherwise we excluded the cell line. We focused on the target pairs rather than specific drug pairs, in order to derive more robust insights.

This leaves us with the following 7 target pairs: AKT/EGFR, AKT/MTOR, BCL2/MTOR, EGFR/MTOR, AKT/BCL2, AKT/ALK and AKT/PARP1, each representing several distinct drug combinations (3, 5, 3, 4, 4, 6 and 4, respectively). We applied our methodology on those target pairs (**Methods, Supplementary text 1**, obtaining the following prediction performances (Pearson’s correlation of observed versus predicted synergy scores): AKT/EGFR (r=0.15), AKT/MTOR (r=0.086), BCL2/MTOR (r=0.2), EGFR/MTOR (r=0.43), AKT/BCL2 (r=0.19), AKT/ALK (r=0.33) and AKT/PARP1 (r=0.50) (**Supplementary Table 1, Fig. 3**), with an average performance of r=0.27 using Leave One Out Cross Validation (**Methods**, **Supplementary Table 1**).

**Fig. 3:**
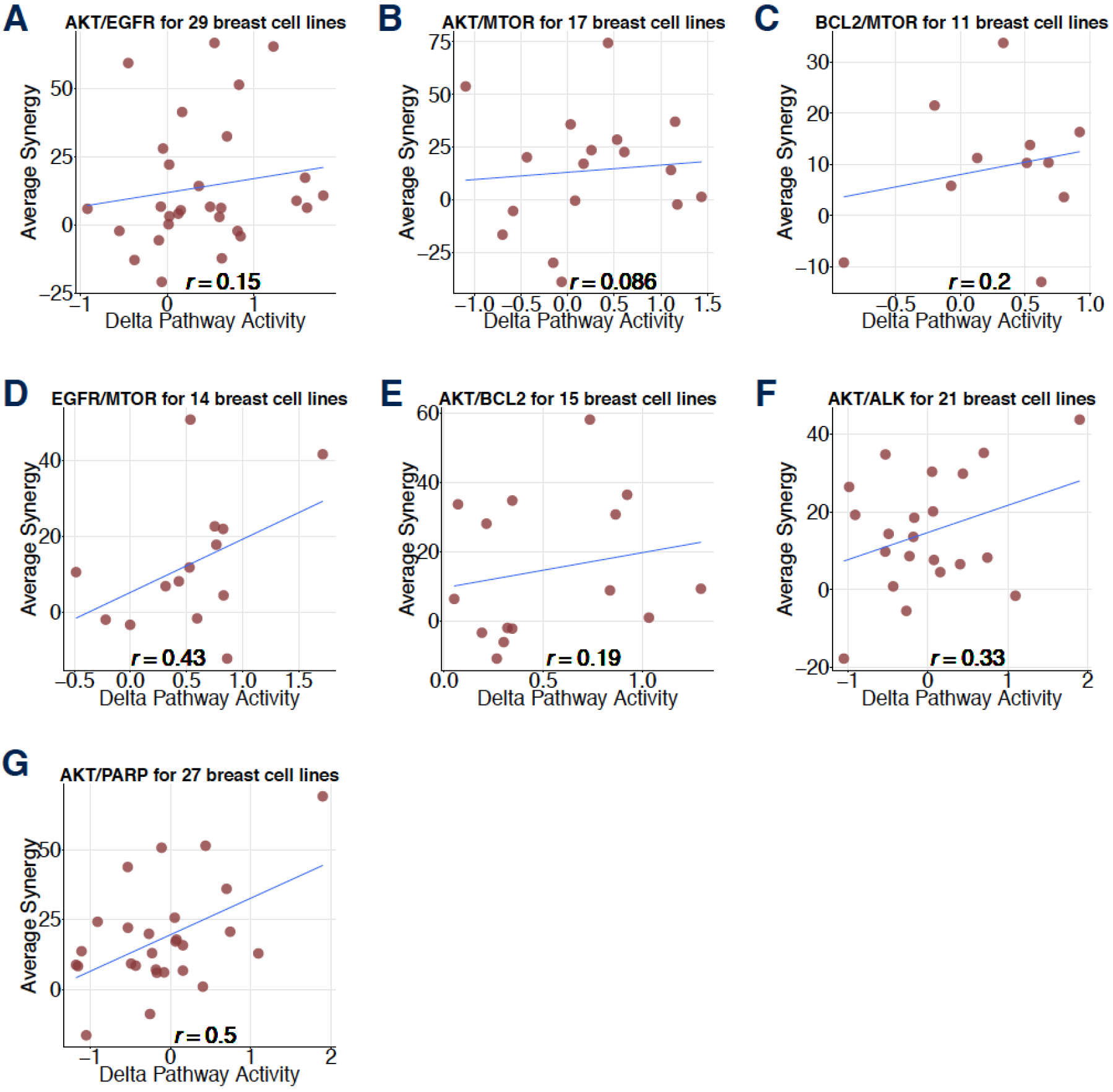
Prediction of drug synergy on breast tissue. **A**, **B**, **C**, **D**, **E, F** and **G** show the prediction result for AKT/EGFR, AKT/MTOR, BCL2/MTOR, EGFR/MTOR, AKT/BCL2, AKT/ALK and AKT/PARP1 targets pairs on breast tissue, respectively (from AstraZeneca DREAM challenge data). Each dot represents a cell line; y axis is the average observed synergy score of the top three drug1-drug2-cell line triplets; x axis is the predicted synergy score (Delta Pathway Activity).

Overall, among all target pairs, AKT/EGFR, AKT/MTOR, BCL2/MTOR, EGFR/MTOR and AKT/BCL2 were predicted with Model 1 (synergy by similarity). AKT/ALK and AKT/PARP1 were predicted using Model 2 (synergy by dissimilarity).

#### Independent experimental validation on colorectal cancer cell lines

We then attempted to experimentally validate our synergy stratification workflow in colorectal cancer. In order to ascertain our method’s capability to detect synergy, we chose a drug combination in the following way:

i. We focused on drug combinations involving the protein BRAF in colorectal cancer, which is frequently mutated in this cancer type (~10% of The Cancer Genome Atlas (TCGA) patients(19)) and can result in uncontrolled, non–EGFR-dependent cellular proliferation(20). Despite the success of BRAF inhibitors in melanoma, in colorectal cancer BRAF monotherapies largely fail to demonstrate clinical efficacy in BRAF^V600^-mutants. BRAF^V600^ is the most common mutation for BRAF (90% cases) where valine is substituted by glutamate in the codon 600. Such mutation can lead to a 500-fold increased activation, stimulating the constitutive activation of MEK/ERK pathway in tumor cells(21). Thus, there is a need for novel combination therapies(22–24) with BRAF inhibitors in colorectal cancer.
ii. We exclude combinations that target, on average, more than three proteins, since too many targets can render the model less precise. There are 101 targets in GDSC besides BRAF, and we computed their functional similarities with BRAF. Insulin Receptor (IR) ranked first with a target functional similarity of 0.73 for the BRAF/IR pair. Insulin has been described as promoting cell proliferation in colorectal cancer by activating MAPK signaling(25), which could explain a similarity with BRAF. Therefore, we chose BRAF/IR as a candidate for validation. We used Dabrafenib as a BRAF inhibitor and BMS−754807 as a selective inhibitor of IR (26).
iii. Since BMS−754807 also targets Insulin Growth Factor 1 Receptor (IGF1R), we chose the proteins BRAF, IR, and IGF1R as drug targets and derived the Delta PA to predict synergy (**Methods, Supplementary Fig. 4h**, **Supplementary Table 1**).

We experimentally validated our methodology on newly generated combination screenings with Dabrafenib and BMS-754807 on 48 colorectal cancer cell lines from the GDSC panel. The synergy score is computed with DeltaXMID (**Methods**). The Pearson’s correlation of observed versus predicted synergy score is 0.31 for all 48 cell lines (**Methods**, **Supplementary Table 1, Fig. 4a**).

**Fig. 4:**
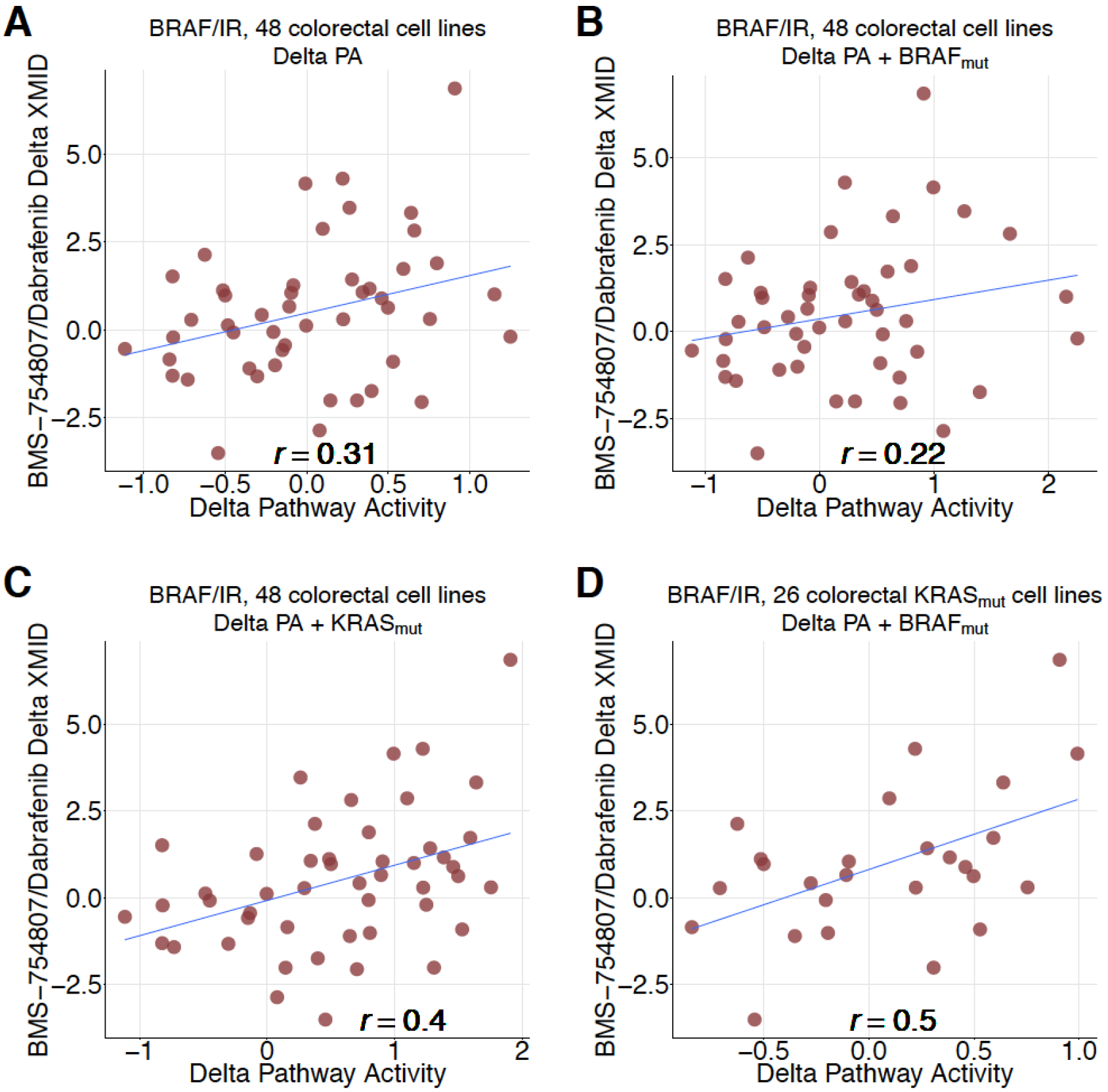
Prediction of BRAF/IR synergy on colorectal tissue. **A** prediction result of BRAF/IR (BMS−754807/Dabrafenib) on all 48 colorectal cancer cell lines. **B** result with BRAF status included in Delta PA formula. **C** result with KRAS status included in Delta PA formula. **D** result on the subset of 26 KRAS_mut_ colorectal cancer cell lines with BRAF status included in Delta PA formula.

We further reasoned that inclusion of information about top predictive pathways should increase the predictive power. In the case of the Dabrafenib and BMS-754807, the most predictive pathway is Hypoxia (**Methods**). KRAS mutation has been shown to differentially regulate the hypoxic induction of HIF-1α and HIF-2α in colon cancer(27). Hence, we added KRAS status in the Delta PA formula and the prediction performance rose to 0.4 (**Fig. 4c**). Accordingly, adding BRAF mutation did not improve the performance (**Fig. 4b**), as MAPK was not as highly ranked as Hypoxia pathway for this combination. However, the performance rose to 0.5 by including BRAF status in the Delta PA formula for the subset of 26 KRAS mutant cell lines (**Fig. 4d**), suggesting an effect of BRAF mutations only when KRAS is mutated. This can have clinical relevance for the around 1% of TCGA colorectal cancer patients have both KRAS and BRAF mutated(19).

### Comparison with supervised learning approaches

Supervised learning has long dominated the prediction of drug pharmacological effect. However, it’s intrinsic reliance on training data makes it difficult to predict in a data-sparse situation. Furthermore, the prediction itself generally does not bring any biological insights (**Supplementary Table 2**). Having these differences in mind, we performed a comparison with our hypothesis based method for synergy stratification.

In the AstraZeneca DREAM challenge, an ensemble of best performing models was trained on the AstraZeneca DREAM combinatorial data, and consecutively tested on an independent combinatorial screen from Merck(28), which achieved a weighted mean correlation of 0.15-0.17. In comparison, we used our synergy stratification workflow on the GDSC panel for hypothesis generation and the AstraZeneca dataset for testing, considered the setting of predicting synergy of new drugs on new cell lines (**Supplementary Fig. 5**). For the seven target pairs (29 drug combinations) from breast tissue, we were able to reach an average drug-wise correlation of 0.27. Of note, the two methods are of very different nature and have very different applications. Therefore, prediction performances should not be compared directly. In the DREAM challenge, synergy scores of drugs/samples are predicted without any prior knowledge of whether a drug combination leads to synergy or not. In contrast, in our synergy stratification workflow, we assume that at least one drugs/samples is synergistic and consecutively predict the stratification based on pathway activity and mutational profiles.

## DISCUSSION

In this paper, we presented an approach with two workflows, one for drug synergy prediction and one for synergy stratification. The synergy prediction workflow can be a powerful framework for compound prioritization in large scale drug screenings. For instance, only testing drugs targeting two functionally very similar or very opposite proteins (|correlation| > 0.7) could significantly reduce the search space, therefore decreasing the cost of drug combination screens. We validated our result on the NCI-ALMANAC dataset(17). Our other workflow for synergy could potentially be used to maximize the drug efficacy of drugs already known to induce synergy, by choosing on which cells (and eventually patients) to apply them. Indeed, knowing that a pair of compounds can be synergistic does not tell us on which cases we should to use it. As real world use case, we envision that for any drug combination described as synergistic, this method could potentially inform about the subset of patients most likely to benefit, based on their transcriptomics profiles, provided enough good quality data to apply the methods.

We introduced the notion of functional similarity between two target proteins. This metric shed lights on two scenarios where drug synergy occurs: when drugs are targeting functionally similar proteins (AKT/EGFR, AKT/MTOR, BCL2/MTOR, EGFR/MTOR and AKT/BCL2) and when they are targeting functionally opposite proteins (AKT/ALK and AKT/PARP1). We hypothesize that combinations of functionally similar targets may lead to synergy due to inhibition of compensatory feedback loops and/or increases target inhibition, whilst functionally dissimilar targets are more likely to be synergistic by targeting potential escape mechanisms. Our results support that synergy occurs and is much easier predicted when the targets are functionally very similar or very opposite. Portraying the interaction between target proteins and pathway activities allowed us to recognize the different synergy cases. Based on that, we applied our method to seven target pairs (AKT/EGFR, AKT/MTOR, BCL2/MTOR, EGFR/MTOR, AKT/BCL2, AKT/ALK and AKT/PARP1, for 29 drug combinations) in breast cancer cell lines. Finally, we predicted and validated a drug combination synergy (Dabrafenib/BMS−754807) on 48 colorectal cancer cell lines.

There are several limitations to this study that can be the focus of future work: (i) The synergy models are simple and could be extended to take into account non-linear effects of pathways adding coefficients to each pathway and including logic (AND/OR) gates. But this would require an extensive training set. (ii) In this present work, target functional similarity is defined with respect to 11 PROGENy pathways, which do not necessarily capture all cancer mechanisms. Besides, only a few cell lines have been used for perturbation experiments to represent each tissue, which does not necessarily capture the whole complexity of the cancer specific signaling mechanism. Expanding this geneset to include more pathways, as well as using more cell lines for each cancer type is likely to improve our models. (iii) In order to predict synergy of new compounds, drug targets have to be profiled by large scale monotherapy drug screening experiments across hundreds of cell lines. Thus, to increase the space of combinations, we require the corresponding monotherapy data. This cost provide important gains, as cost for monotherapy grows linearly, but exponentially for combinations. (iv) There is currently limited tissue specific publicly available drug synergy data. Such datasets could be highly valuable to further refine our and other approaches.

Our study findings are aligned to those of the DREAM drug combination challenge(7), where synergy was found to be highly context dependent. In our case, we predicted synergy with a linear combination of pathway activities. Bansal et al. predicted synergy from single-compound perturbation data. They found that synergy occurs for drug pairs which induce very similar or very opposite gene perturbation statuses. We used single-compound drug response data and the Macau algorithm to compute the target functional similarity, which reflects the similarity of drug response changes for different pathways after targeting a specific protein. We found that compounds that have very similar or very opposite functional profiles tend to be more synergistic. We used the inferred synergy mechanism and pathway activities to predict synergy of new compounds.

We compared our synergy stratification workflow with state-of-the-art supervised learning (**Supplementary text 2, Supplementary Table 2**) and highlighted the pros and cons for each methodology: (i) Naive supervised learning approaches are easy to implement, do not require extensive domain expertise (although still highly valuable), and can be used for all possible prediction settings (**Supplementary Fig. 5**). However, they require an extensive set of drug combination drug response data as training set. (ii) For our synergy stratification methodology, linear combination of pathway activities is well suited for biological interpretation. However, it can only be used in drug-wise setting and requires significant domain knowledge and literature evidence.

In terms of translatability to the *in vivo* setting, supervised learning needs combination data of the organism of interest for training, whereas our method requires the target functional similarity to be built based on monotherapy data on the organism of interest. We attempted to test our methodology to Novartis PDX data(31) but it was not possible due to the fact that the drug targets and gene expression were not predictive of the single drug response, with prediction performance at random level. Therefore, we couldn’t derive a robust and reliable interaction matrix/Target functional similarity from this data(15). The random performance of single drug response prediction on Novartis PDX data(31) might be explained by (i) the high level of missing values of the drug response matrix (64%), (ii) the model system could be so complex that transcriptomics alone does not reflect the underlying biology enough to be predictive of drug response, and (iii) that measurements could have been inaccurate, owing to a lack of standardization for this new technology. When larger and more mature datasets of this kind become available, it will be possible to test our approach on such an *in vivo* context.

Palmer and Sorger stated that successful drug combination in tumour shrinkage are mostly due to targeting unrelated pathways, without any real synergy(30). They define drug action similarity by the correlation of single drug response data, which resembles our use of target-pathway based similarity score. They concluded that highly correlated independent drug responses can explain the majority of combination clinical trial (synergy), whereas lowly correlated independent drug responses makes independent action of drugs the dominant mechanism in clinical populations (additivity). While the analysis is different as we used cell line data, there are commonalities in our findings.

In summary, exploring the interactions between drug targets and signaling pathways in a tissue specific manner can provide a novel in-depth view of cellular mechanisms and drug modes of action, which can ultimately rationalize drug combination strategies in cancer. Target functional similarity could be used as a metric for compound prioritization. Synergy by similarity hypothesis could be a rational for first line treatment, while synergy by opposite effect could potentially fit patients having acquired resistance.

## METHODS

### Matrix factorization with Macau

Macau trains a Bayesian model for collaborative filtering by also incorporating side information on rows and/or columns to improve the accuracy of the predictions(14) (**Fig. 1a**). Drug response matrix (IC50) can be predicted using side information from both drugs and cell lines. We use target protein as drug side information and transcriptomics/pathway as cell line side information. Each side information matrix is then transformed into a matrix of L latent dimension by a link matrix. Drug response is then computed by a matrix multiplication of the two latent matrices. Macau employs Gibbs sampling to sample both the latent vectors and the link matrix, which connects the side information to the latent vectors. It supports high dimensional side information (e.g. millions of features) by using conjugate gradient based noise injection sampler.

### Pathway activities

We transformed the transcriptomics data into pathway activity scores using PROGENy (**Fig. 1a**). PROGENy is a data driven pathway method aiming at summarizing high dimensional transcriptomics data into a small set of pathway activities. PROGENy derives pathway signatures from the genes that are altered when perturbing a pathway instead of solely from the genes within the pathway as other methods do. This improves the estimation of pathway activities(12). We currently use the following 11 PROGENy pathways: EGFR, NFkB, TGFb, MAPK, p53, TNFa, PI3K, VEGF, Hypoxia, Trail and JAK-STAT.

### Interactions between drug target and pathway activities

We computed the interactions between target proteins and signaling pathway activation status with respect to drug response (IC50) using matrix factorization (**Fig. 1a**). This interaction can be defined as the importance for those two entities to be simultaneously involved in order to have an impact on drug response(15), e.g. how the simultaneous activation of a certain pathway and targeting a certain protein can be associated with drug response. For instance, a strong interaction between protein MEK1/MEK2 and pathway EGFR in pancreatic cancer is interpreted as follows: Activation of the EGFR pathway correlates with sensitivity when targeting MEK1/MEK2. If this were a causal relationship, it could mean that EGFR pathway activation confers sensitivity to any drug targeting protein MEK1/MEK2.

We used the GDSC cell line panel that contains drug response (IC50) data of 265 drugs on 990 cell lines. For each of the 16 tissues (with more than 20 cell lines), we computed the interaction matrix between drug targets and pathway activities using the multitask learning algorithm Macau(14,15). Our algorithm tries to learn multiple tasks (predicting multiple drugs) simultaneously and uncovers the common (latent) features that can benefit each individual learning task(29). We used manually curated target proteins for the drug (**Supplementary Table 4**), and gene expression derived pathway scores for the cell lines. The interaction matrix gives hints about the drug’s mode of action, by uncovering in which condition (pathway status) targeting a certain protein correlates with higher drug sensitivity.

### Building specific models for each synergy hypothesis

The core idea of the synergy stratification workflow is to predict the synergy score using the pathway context in addition to drug target information. We believe that synergy is not only due to the drugs’ properties, but also dependant on the signaling context.

#### Synergy Model 1 (Maximizing drug sensitivity)

For functionally similar target pairs (Mechanism 1), we rank the pathways based on their sensitive or resistant interaction profile with respect to the drug targets. We postulate that synergy is maximized under a pathway condition where both drugs’ effects are maximized. The optimal condition for synergy is therefore when pathways associated with drug sensitivity are upregulated, and pathways associated with drug resistance are downregulated (**Supplementary Fig. 3**). As a consequence, if two target proteins have strong functional similarity e.g. high correlation between their interaction profile with pathway activities, synergy is maximized when the sensitizing pathways are activated and pathways conferring resistance not activated. We predict synergy by taking the average of the top N sensitive pathway scores, subtracted by the average of the top M resistant pathway scores. Therefore, for each cell line, we introduce the concept of Delta Pathway Activity (Delta PA) to predict synergy:

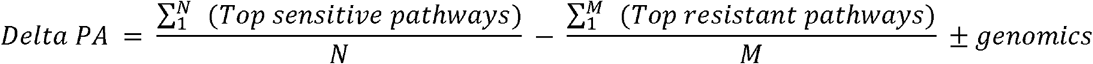

We compute the average pathway score for both sensitive and resistant groups. Each group should include a minimum of one to a maximum of three pathways. We select the top pathways with group membership thresholds determined by cross validation. If applicable, we include in the formula the genomic information which can be mutation (SNP) or copy number variation (CNV). For instance if protein EGFR is targeted, we include CNV_*EGFR*_. Group membership parameters are defined using cross validation.

### Synergy Model 2 (Maximizing drug resistance)

For functionally opposite target pairs (Mechanism 2), when a pathway’s activation is associated with resistance for one target protein, it is also associated with sensitivity for the other target protein, to compensate. Two drugs can be individually ineffective, but more effective when combined. Therefore, synergy may arise in a situation of drug resistance. This could be explained by the fact that if a cell line is resistant for one (or both) of the drugs, there is “more opportunities” to be synergistic. When both drugs kill a given cell very efficiently, there is no synergy, as both drug A alone, drug B alone and combination A + B can kill all the cells. Unsurprisingly, resistance biomarkers were found to be predictive of synergy in the recent AstraZeneca DREAM challenge(10). Therefore Delta PA should maximize the pathways conferring resistance and minimize the sensitizing pathways. The formula becomes:

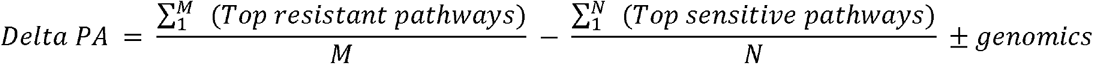

Model 2 is less likely to suit functionally similar pairs (Mechanism 1). If the two drugs have similar functional profile, maximizing the resistance scenario equals increasing the dose of the same inefficient drug, thus, unlikely to improve the outcome. Likewise, Model 1 is less suitable for Mechanism 2, as maximizing the sensitizing pathways is the same as prioritizing a situation where drug 1’s sensitive effect outweighs drug 2’s resistant effect. Thus, Mechanism 2’s core idea would become obsolete, as by definition, the resistance scenario must prevail in case of escape mechanism. Of note, having an opposite functional profile does not imply Mechanism 2. An opposite pathway-response profile for two targets, offers the “functional scenario” for the cell to escape the damage induced by one drug. Yet, there could still be a scenario which maximizes the sensitizing pathways. This corresponds to two drugs targeting completely independent pathways, which is more due to independent actions rather than additivity or synergy(30).

### General framework for predicting synergy score

**Step 1**: For two given target proteins T1 and T2, find their interactions with the PROGENy pathways using Macau (**Supplementary Fig. 4**).

**Step 2A**: If available, use literature to guide the choice of Model e.g if we know that a drug combination is synergistic when a pathway X is activated, the model would be the one which gives a positive sign for pathway X. Otherwise go to **Step 2B**.

**Step 2B**: Compute the functional similarity between T1 and T2 (Pearson’s correlation between T1 and T2’s interactions with the pathways).

If the correlation is close to 1, use **Model 1** to define the Delta PA formula.

If the correlation is close to −1, use **Model 2** to define the Delta PA formula.

If the correlation is between −0.4 and 0.4, it is an undetermined case.

**Step 3**: Find top sensitive and top resistant pathways (as previously described in synergy models). Take into account literature evidence in choice of pathways (for known drugs or targets). If a pathway is described as important in literature but does not appear in top three of a group, we include it, as well as any pathway separating the first from the one of interest, while respecting the limit of three pathways per group.

**Step 4**: In case of multiple drugs representing the same target pair, as in the AstraZeneca data set, keep the drugs that are specifically targeting the two proteins of interests, while removing those having off target effects (at least 3 drug pairs left).

**Step 5**: Use the Delta PA formula to predict synergy of a drug combination targeting T1 and T2. The pathway activities of the formula are computed by PROGENy on the cell lines of interest.

More details about the cross validation procedure and synergy score computation are in **Supplementary Methods.**

The source code to generate the interaction matrices is available at: https://github.com/saezlab/Macau_project_1

The source code of the result analysis is available at: https://github.com/saezlab/Macau_Synergy_Prediction

GDSC data were downloaded from: http://www.cancerrxgene.org/

Drug IC50 version 17a
Basal gene expression 12/06/2013 version 2
Drug target version March 2017

DREAM drug combination challenge data were acquired through an AstraZeneca Open Innovation Proposal.

NCI-ALMANAC data is downloaded from the publication Holbeck et al.(17)

Novartis PDX data is downloaded from the publication Gao et al.(31)

## ACKNOWLEDGEMENTS

We would like to thank Federica Eduati and Bence Szalai for their suggestions and ideas in improving the manuscript, as well as Francesco Ceccarelli and Javier Perales Patón for checking code reproducibility.

## SUPPLEMENTARY DATA

### Supplementary Methods

#### Cross validation of group membership thresholds

##### Prediction using a fixed threshold for both groups

For computing the Delta Pathway Activity formula, we chose the group membership to be the same in the top sensitive and top resistant groups. We only included pathways producing more than 70% of the effect of the first selected pathway. This choice of parameter seems to be a reasonable choice between including no additional pathway (threshold close to 100%) and including too many potentially irrelevant pathways (50%). We did a sensitivity analysis for this parameter and used different thresholds for the two different groups (sensitive and resistant). We fixed one group’s threshold while varying the other group’s threshold. We observe that for many target pairs, the sensitive group’s threshold is rather stable (**Supplementary Fig. 6**). For the sensitive group, the model is quite robust to variation of this parameter, whereas for the resistant group, a lower threshold (including more pathways) seems to yield better result. This could be explained by the fact that GDSC drug screening adjusts the drug concentration so that only a few cell lines respond while most are resistants. Therefore, more information are reflected from the resistant side than from the sensitive side.

##### Prediction using cross validated and different thresholds for each group

We next predicted each target pairs of AstraZeneca breast data with a Leave One Out Cross Validation (LOOCV) to optimize the group membership thresholds. For each target pair, we used as thresholds the best values all target pair of the training set. The average prediction of the 7 target pairs is 0.27. Finally, we used the average parameters for the breast data to predict the BRAF/IR in colon data (**Supplementary Table 1**).

#### Drug synergy metrics

For AstraZeneca dataset, drug effects on cancer cell lines are measured at several concentrations for each drug in a 5-by-5 matrix format. Therefore, the effect is described by a dose-response surface rather than a curve. The benefit of a drug combination can be partly assessed by the extra effect obtained when combining the drugs. Drug combinations are classified as synergistic, additive or antagonistic, based on the deviation of the observed drug combination response from the theoretical additive response. The theoretical additive response is quantified with the Loewe additivity model(32–35) based on the monotherapies of both drugs. Loewe additivity assumes the two drugs act on a protein through a similar mechanism. Synergy score is quantified with Combenefit(18), which calculates the volume between the experimentally observed on theoretical response surface. As in the AstraZeneca DREAM challenge, we consider a score greater than 20 as synergistic(10).

In colorectal cancer, we tested the drug combination of BMS-754807 and dabrafenib in 48 colorectal cancer cell lines. BMS-754807 (S2807, Selleckchem) was screened at 0.5 μM against a 7 point dose response of dabrafenib (S1124, Selleckchem), ranging from 10 nM-10 μM. The XMID, which is akin to an IC50, of dabrafenib alone and dabrafenib in combination with BMS-754807 were calculated and the ΔXMID=XMID(dabrafenib)-XMID(dabrafenib+BMS-754807) calculated. The fold difference in XMID can be calculated by y-fold=2^ΔXMID, hence a ΔXMID of 3.32 corresponds to a 10-fold lower XMID for dabrafenib + BMS-754807 compared to dabrafenib alone.

### Supplementary text: Methodology applied to breast tissue

We explained through target-pathway interactions, two mechanisms of drug synergy. In order to validate our synergy models, we first looked at public data, using the DREAM AstraZeneca drug combination challenge(10), which experimentally tested >120 folds drug combinations compared to the previous Bansal et al. challenge. Furthermore, the AstraZeneca challenge expanded the number of tested cell lines including their deep molecular characterisation enabling for the first time identification of synergy biomarkers. We tested our model on 7 target pairs (29 drug combinations) from the AstraZeneca DREAM challenge(10), and chose breast as the most represented tissue with 33 cell lines.

We applied our general framework to predict synergy scores. The first step was to determine the top sensitive and top resistant pathways for a certain target - pathway pair (**Supplementary Fig. 4**). We then derived the formula of Delta Pathway Activity and predicted the drug synergy (**Supplementary Table 1**). When choosing between Model 1 and 2 for the synergy model, the target functional similarity was the main criteria. If the similarity is close to 1, we use Model 1. If the similarity is close to −1, we use Model 2.

PI3K/AKT/MTOR pathway plays a significant role in treatment resistance in breast cancer(36). Therefore, we hypothesized that the PI3K pathway will be informative of the synergy if AKT is targeted. Therefore, each time AKT is targeted, we included PI3K pathway as well as any pathway between the first one and PI3K, while respecting the limit of maximum three pathways per group.

When grouping pathways in the top sensitive and top resistant groups, we consider only those that have at least one significant interaction with the drug targets. If not significant, we discard the pathway. Exceptions are made when only one pathway is included (the top sensitive or top resistant one) and when the pathway has a stronger interaction than a pathway included by prior knowledge (literature).

For AKT/EGFR (**Supplementary Fig. 4a, Fig. 3a**): the top sensitive pathway is EGFR and the top resistant are MAPK. The target functional similarity between AKT1/2 and EGFR is 0.9 (**Supplementary Table 1**). Therefore, we used synergy Model 1. Since protein EGFR is targeted, we also added CNV information:

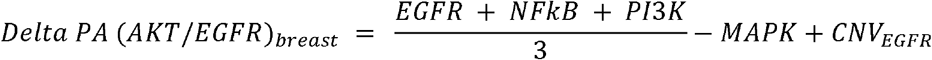
For AKT/MTOR (**Supplementary Fig. 4b, Fig. 3b**): the top sensitive pathways are EGFR and VEGF. The top resistant pathways are MAPK and TNFa. The target functional similarity between AKT1/2 and MTOR is 0.8 (**Supplementary Table 1**). Therefore, we used synergy Model 1:

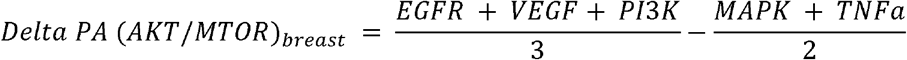
For BCL2/MTOR (**Supplementary Fig. 4c, Fig. 3c**): the top sensitive pathways are VEGF, NFkB and Trail and the top resistant pathways are MAPK and TNFa. The target functional similarity between BCL2 and MTOR is 0.7 (**Supplementary Table 1**). Therefore, we used synergy Model 1:

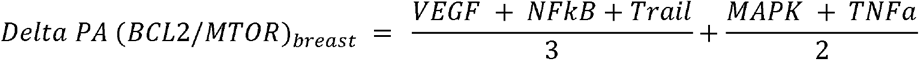
For EGFR/MTOR (**Supplementary Fig. 4d, Fig. 3d**): the top sensitive pathways are EGFR and NFkB. The top resistant are MAPK and TNFa. The target functional similarity between EGFR and MTOR is 0.6 (**Supplementary Table 1**). Therefore, we used synergy Model 1. Since protein EGFR is targeted, we also added CNV information:

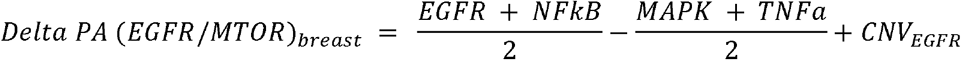
For AKT/BCL2 (**Supplementary Fig. 4e, Fig. 3e**): the top sensitive pathway is EGFR and the top resistant pathway is MAPK. The correlation between AKT1/2 and BCL2 is 0.5 (**Supplementary Table 1**). In this case, we used Model 1:

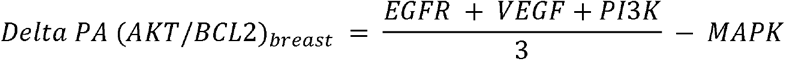
For AKT/ALK (**Supplementary Fig. 4f, Fig. 3f**): the top sensitive pathway is EGFR and the top resistant pathways are MAPK and TNFa. The target functional similarity between AKT1/2 and ALK is −0.4 (**Supplementary Table 1**). Therefore, we used synergy Model 2:

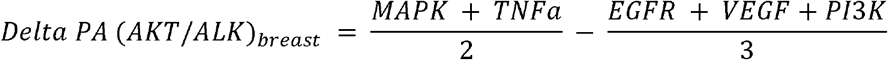
For AKT/PARP1 (**Supplementary Fig. 4g, Fig. 3g**): the top sensitive pathway is EGFR and the top resistant pathways are MAPK and TNFa. The correlation between AKT1/2 and PARP1 is −0.8 (**Supplementary Table 1**). In this case, we used Model 2:

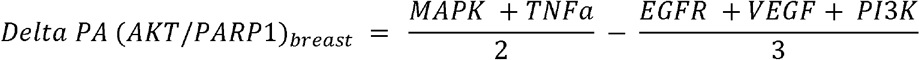

### Supplementary tables

**Supplementary Table 1: Drug synergy prediction for breast and colorectal cancer cell lines.**

**Supplementary Table 2: Comparison of synergy stratification workflow with supervised learning.**

**Supplementary Table 3: Different settings for drug response prediction**

**Supplementary Table 4**: drug target information downloaded from https://www.cancerrxgene.org on March 2017.

### Supplementary figures

**Supplementary Fig. 1:**
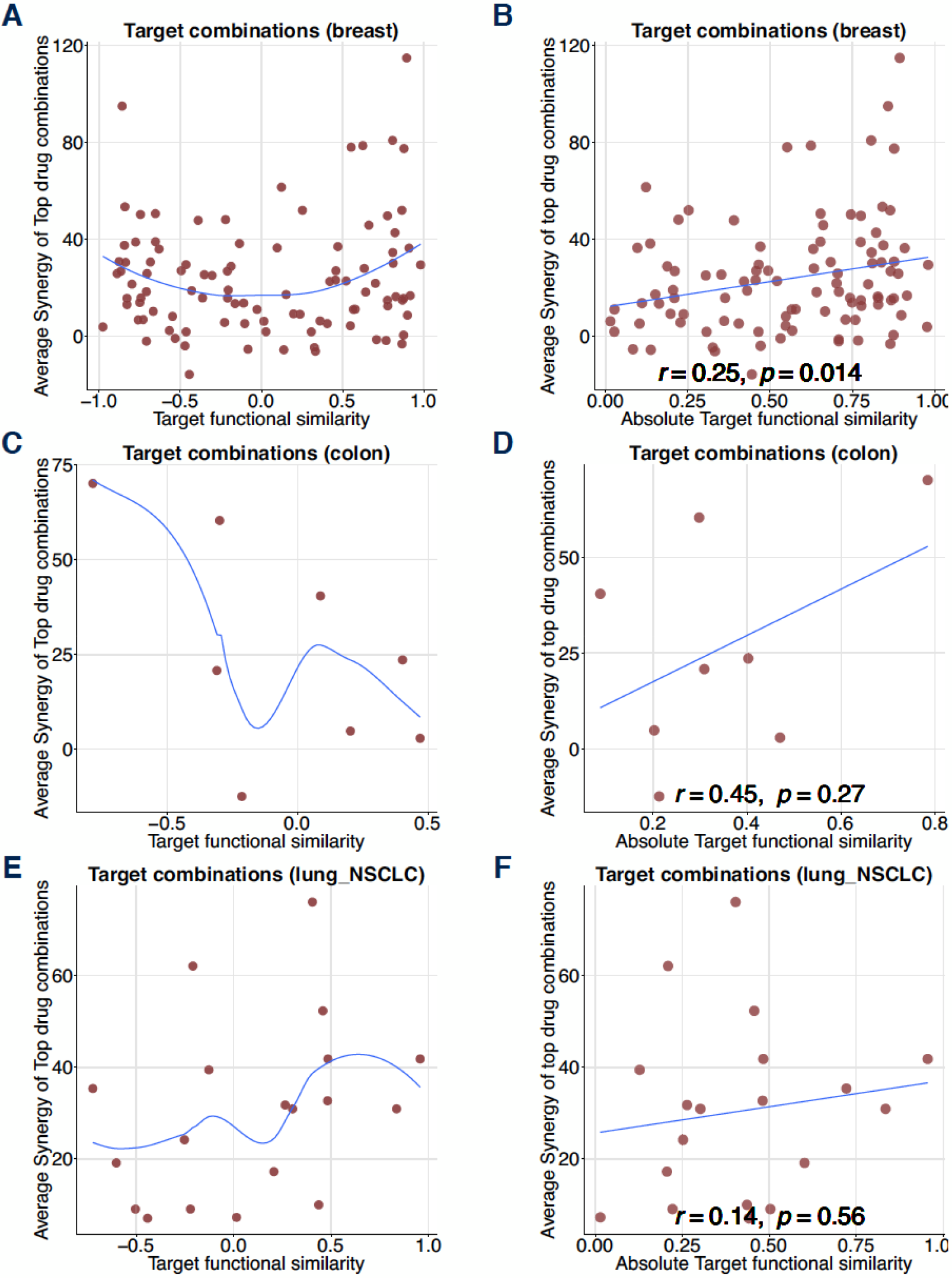
Influence of target functional similarity in drug synergy for AstraZeneca DREAM dataset. The target functional similarity is the correlation between two target proteins by their interactions with the PROGENy pathways. For each tissue, we plot the synergy against the target functional similarity and its absolute value. **A** and **B** for breast tissue. **C** and **D** for colon tissue. **E** and **F** for NSCLC lung tissue.

**Supplementary Fig. 2:**
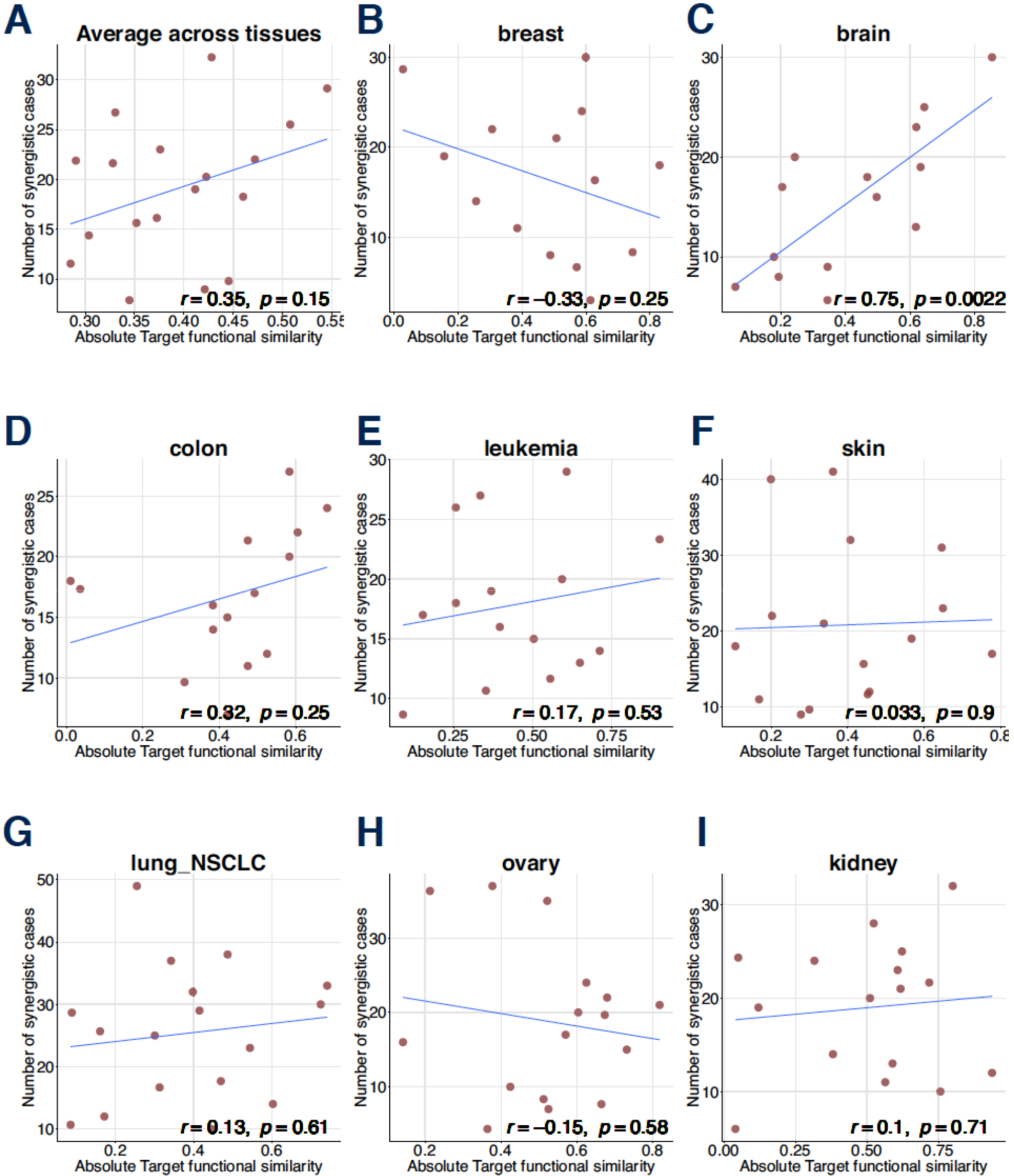
Influence of target functional similarity in drug synergy for NCI-ALMANAC dataset. The target functional similarity is the correlation between two target proteins by their interactions with the PROGENy pathways. For each tissue, we plot the number of synergistic combinations against the absolute target functional similarity.

**Supplementary Fig. 3:**
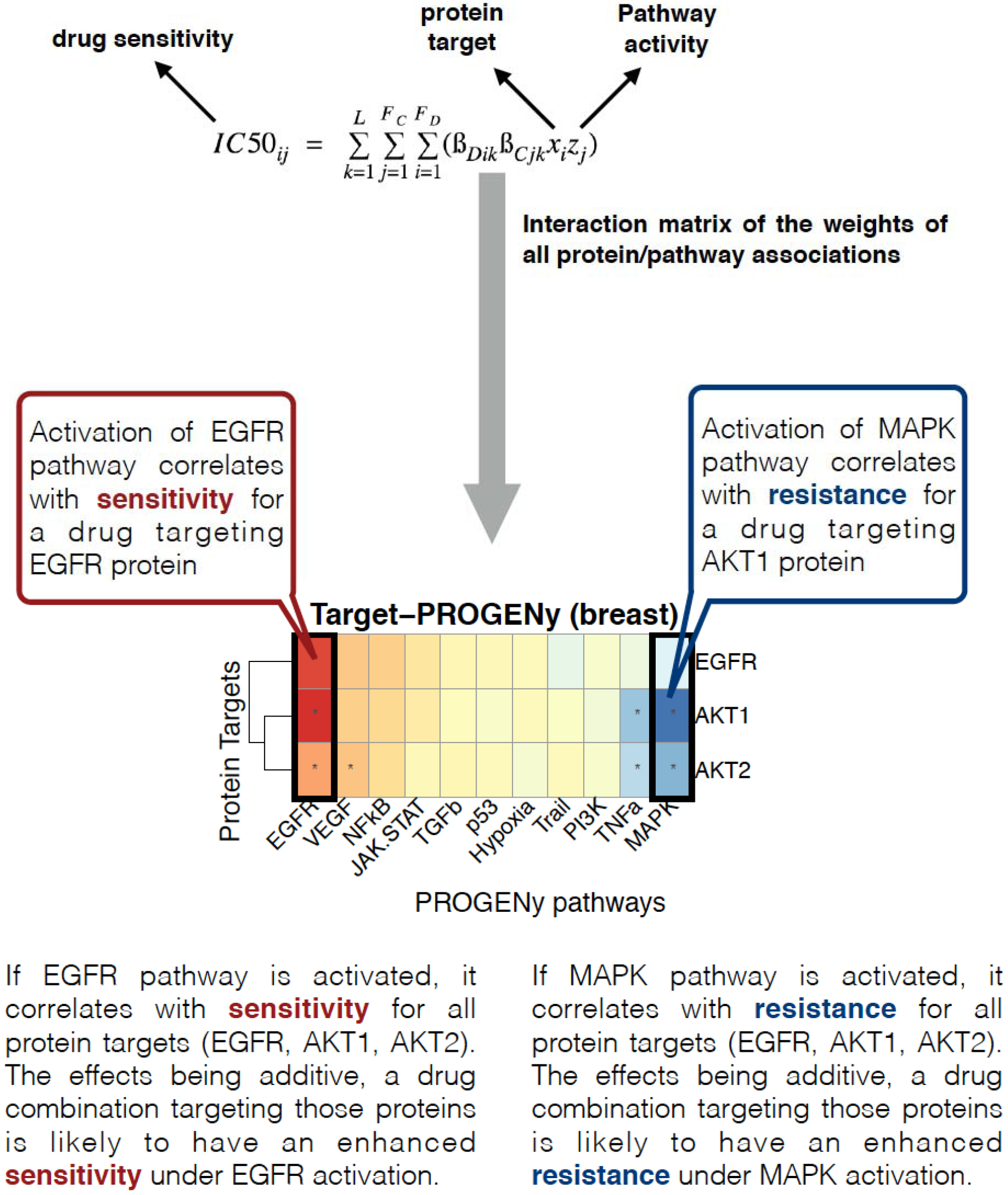
Interpretation of the interaction matrix. Enhanced sensitivity occurs when targeting several proteins involved in drug response under the activation of the right pathway. The same rule applies to resistance.

**Supplementary Fig. 4:**
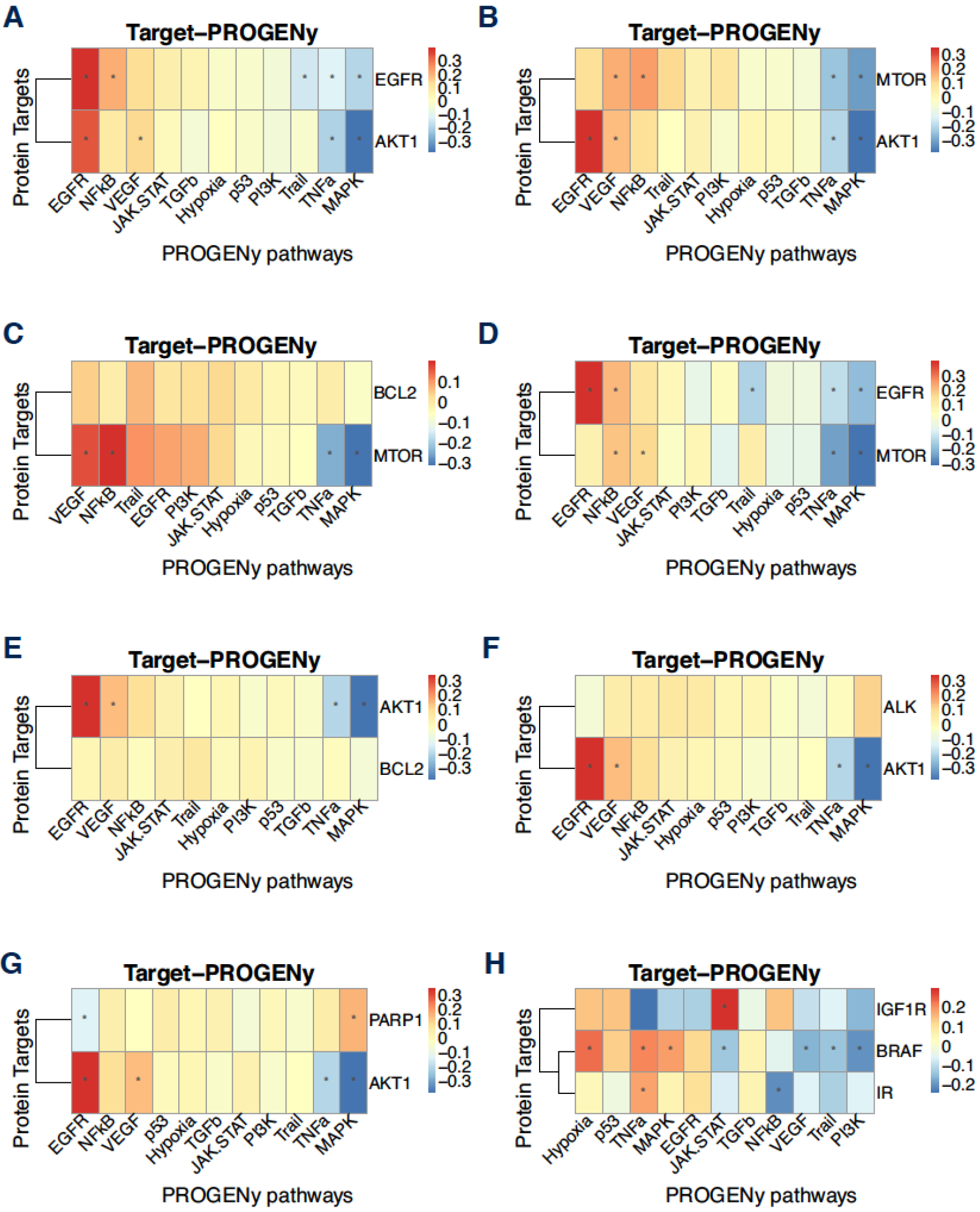
Functional profile of target proteins in breast and colorectal tissues. **A**, **B**, **C**, **D**, **E, F** and **G** describe the functional profile of AKT/EGFR, AKT/MTOR, BCL2/MTOR, EGFR/MTOR, AKT/BCL2, AKT/ALK and AKT/PARP1 pairs in breast tissue. **(h)** describes BRAF/IR’s functional profile in colorectal tissue. The functional profile is a subset of the target pathway interaction in the Macau model. Pathways are ordered from the most sensitizing to the least. Significance of the interaction values is corrected according to Benjamini & Yekutieli procedure (40% FDR) as described in Yang *et al*.(15).

**Supplementary Fig. 5:**
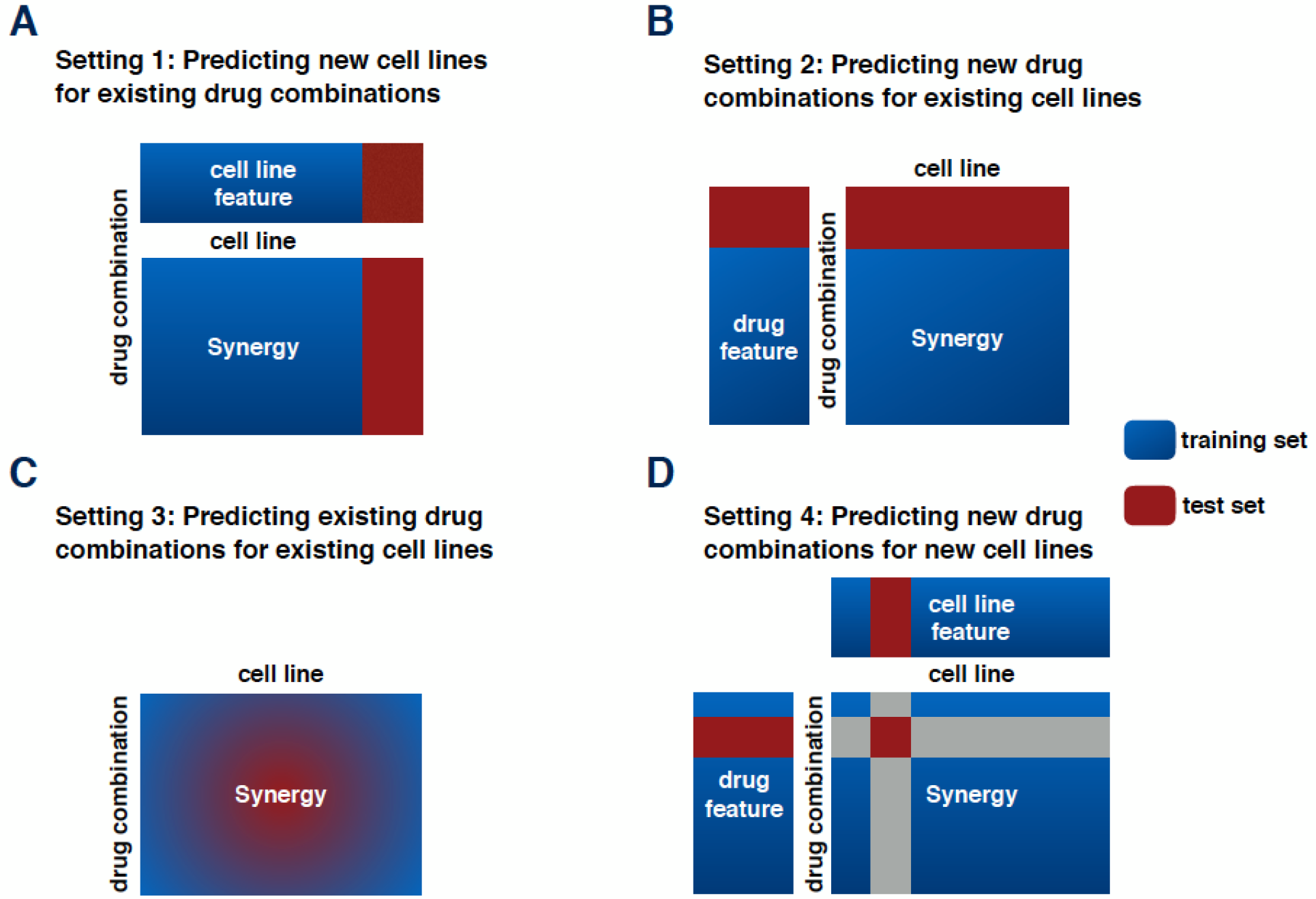
Different settings in drug synergy prediction. **A.** Predicting new cell lines for existing drugs. For each drug pair, we compute the Pearson’s correlation of observed versus predicted synergy across all cell lines of the test set. **B.** Predicting new drug synergy for existing cell lines. For each cell line we compute the Pearson’s correlation of observed versus predicted synergy across all drug pairs of the test set. **C.** Predicting existing drug synergy for existing cell lines. This is a missing value imputation setting where side information of drug and cell lines are not required, but can be used to improve the result. The test data is defined by a percentage of the whole data set. We compute the Pearson’s correlation of observed versus predicted synergy for all randomly chosen drugs - cell line triplets of the test set. **D.** Predicting new drug synergy for new cell lines. We do two simultaneous cross validation on both drug and cell line sides. The test data is defined by association of the test set of the drug side with the test set of the cell lines side. We compute the Pearson’s correlation of observed versus predicted synergy for all drug - cell line pairs of the test set.

**Supplementary Fig. 6:**
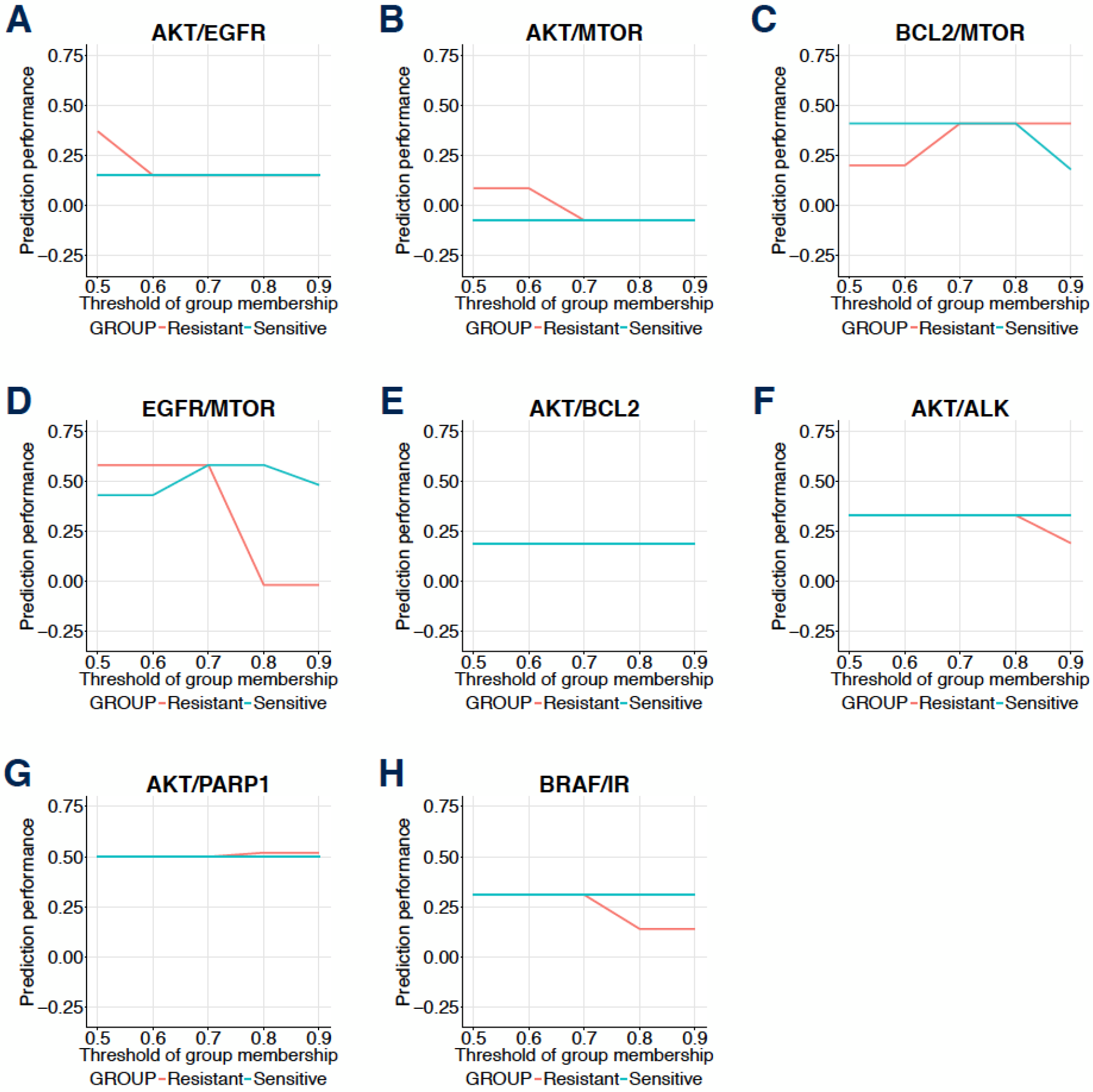
Sensitivity analysis for group membership parameters. In the determination of Delta Pathway Equation, we explored the prediction performance for each target pair in AstraZeneca breast data and colorectal validation data, based on the following parameters: threshold for group membership of the top sensitive pathways and top resistant pathways.

**Supplementary Fig. 7:**
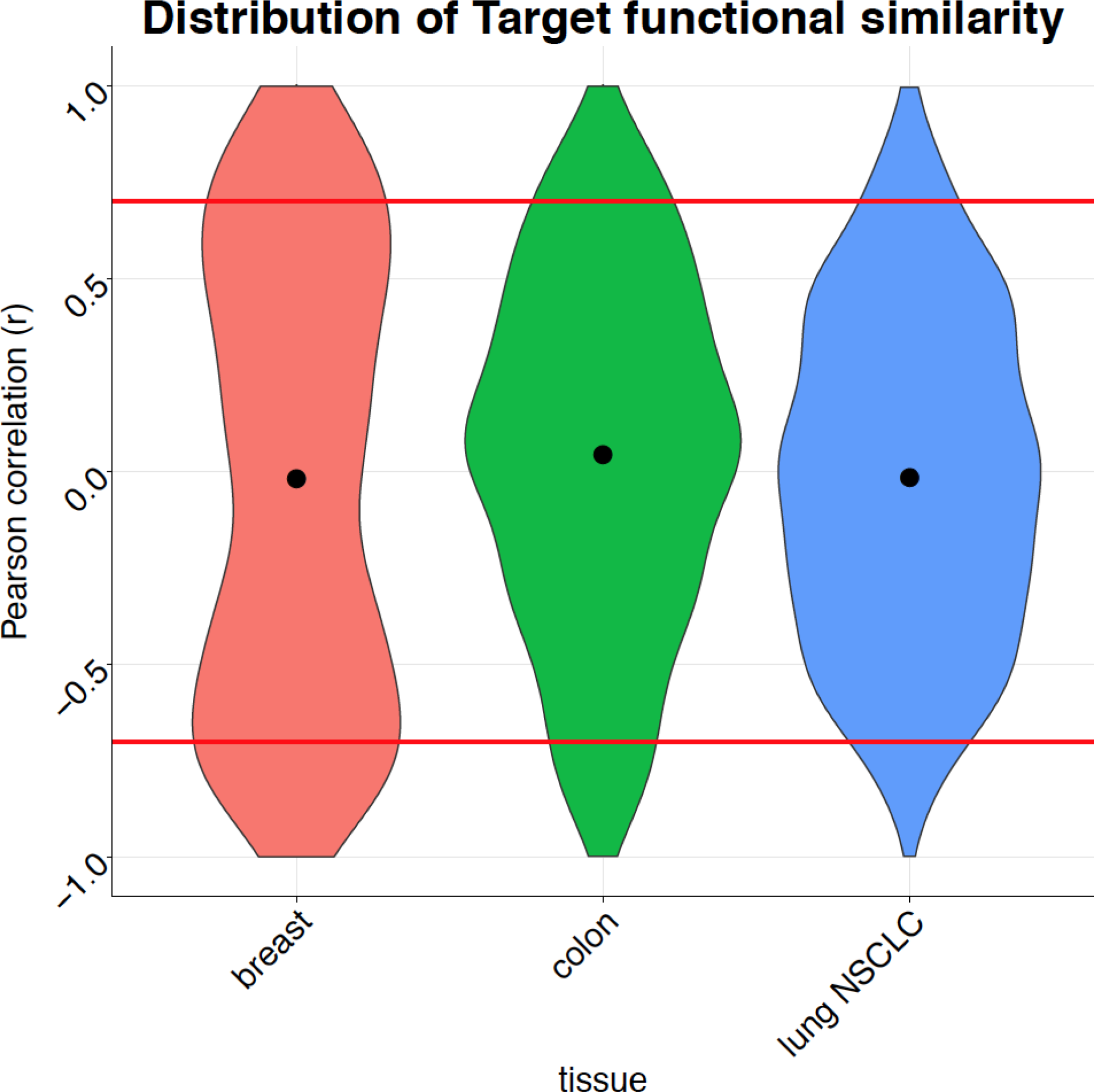
Distribution of similarity values across tissues. For each tissue, we plotted the target functional similarities of the profiled target proteins and set the cut off of high similarity and high dissimilarity at 0.7 and −0.7.

